# Solid-Phase Peptide Capture and Release for Bulk and Single-Molecule Proteomics

**DOI:** 10.1101/2020.01.13.904540

**Authors:** Cecil J Howard, Brendan M. Floyd, Angela M. Bardo, Jagannath Swaminathan, Edward M. Marcotte, Eric V. Anslyn

## Abstract

The field of proteomics has expanded recently with more sensitive techniques for the bulk measurement of peptides as well as single-molecule techniques. One limiting factor for some of these methods is the need for multiple chemical derivatizations and highly pure proteins free of contaminants. We demonstrate a solid-phase capture strategy suitable for the proteolysis, purification, and subsequent chemical modification of peptides. We use this resin on an HEK293T cell lysate and perform one-pot proteolysis, capture, and derivatization to generate a cellular proteome that identified over 40,000 bead-bound peptides. We also show that this capture can be reversed in a traceless manner, such that it is amenable for single-molecule proteomics techniques. With this technique, we perform a fluorescent labeling and C-terminal derivatization on a peptide and subject it to fluorosequencing, demonstrating that washing the resin is sufficient to remove excess dyes and other reagents prior to single-molecule protein sequencing.

## Introduction

With the increasing sensitivity of proteomics methods, a number of new proteins, protein isoforms^[1]^, and post-translational modifications^[3]^ have been discovered. The increase in sensitivity is due to both improvements of the mass spectrometers and the ability to generate high quality protein/peptide samples that are often highly derivatized^[4]^. These methods, however, often utilize purification techniques that are prone to sample losses, and the inclusion of multiple derivatization/purification cycles can lead to low abundance peptides dropping below the detection thresholds^[5]^. This can lead to a bias against rare or low abundance peptides that may be biologically important^[6]^.

The manner that peptides are prepared from biological materials for mass spectrometry analysis is an important consideration in proteomic studies. For example, in bottom-up proteomics, the mode of digestion of proteins is a critical decision. It is routinely done in-solution, where proteases are added to the proteins directly, or the protease treatment is done to specific gel locations after an initial 1D or 2D polyacrylamide electrophoresis separation. After digestion, the sample can be derivatized for several purposes: to eliminate unwanted side products such as disulphides ^[7]^, introduce isotopic labels for quantitation ^[8]^, or to aid ionization ^[9]^, and add handles that can be cleaved to induce specific cleavage patterns ^[10]^. With each of these protocols the preparation requires that the sample be purified to separate the peptides from any side-products or unreacted chemicals.

**Scheme 1.**
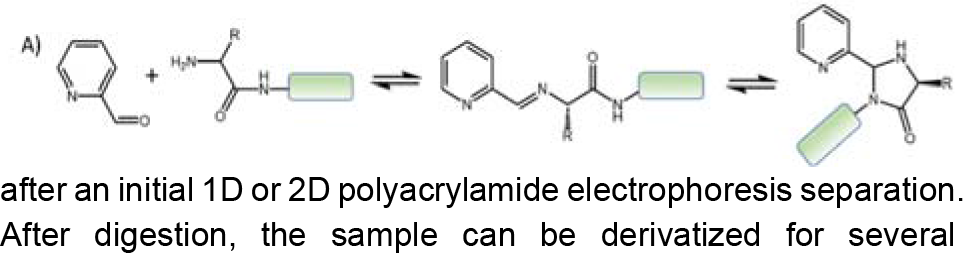
Formation of the N-terminal imidazolidinone cap by pyridinyl carboxaldehyde^[2]^.

One method that we envisioned could be used to improve sample preparation is to bind the proteins/peptides to a bead or other solid matrix. While such an approach has been done before, immobilization typically relies on either the addition of non-natural amino acids as purification handles^[11]^, relies on non-specific hydrophobic precipitation^[12]^, charge based non-covalent bonding, or in some cases metal chelation strategies^[13]^. These properties make them unattractive for studies in which installing non-natural amino acids via amber codon suppression is difficult, such as in mammalian cell cultures^[14]^, or due to the limitations in solvent/buffer conditions that are compatible with non-covalent immobilization techniques,^[13]^ or to the loss of peptides due to non-covalent and weak attachment to the resins.

A method that allows for the binding of peptides on resin support in a covalent and reversible manner would enable complex manipulations with higher overall yields. As demonstrated below in the context of fluorosequencing, the method introduced herein allows for the capture, purification and derivatization of peptides, and in theory, detection of low abundance peptides. Importantly, this procedure allows for derivatization schemes that could otherwise not be performed because of the use of excess reagents and washing steps, analogous to peptide synthesis on resin, where experimental procedures are optimized to impart high yield and speed rather than the need for purification^[15]^.

## Results and Discussion

In order for such a procedure to be applicable to all peptides generated during a proteolytic cleavage, the ideal capture reagent will only react with the N- or C-terminus of a peptide/protein and not with any of the side chains that may be present. A specific reaction with the N-terminus is an attractive choice, as all peptides generated via a protease possess an amine termini, and the protease generally imparts knowledge of either the N- or C-terminal amino acids due to its specificity of cleavage^[16]^. The N-terminus is also a good target, as there is a difference in its ^conjugate acid’s p*K*_a_, as well as nucleophilicity, when comparing^ it to the sidechain amine on lysine^[17]^. While there are methods that allow for the selective modification of the N-terminal amine over lysine amines, most of them rely on permanently modifying the N-terminus, such as creating a new amide bond^[18]^ or converting the N-terminal amino acid into an aldehyde^[19]^. It is also possible to use the C-terminus as a handle for peptide purification, however, due to the similarities between the side chain carboxylic acids of Asp and Glu and the C-terminus this is a less attractive option (the p*K*_a_ of these three range from 3.3-4.2)[20].

Here we describe the use of imidazolinone formation from the N-terminus of peptides using 2-pyridinyl carboxaldehyde (2PCA) (Scheme 1) as first introduced by MacDonald et al. ^[2]^. This was an attractive option for a next-generation capture resin that would have no cross-reactivity with lysine residues, unlike current styrene-aldehyde resins that rely on unstable imine formation with Lys and the N-terminus^[21]^. However, in order to be a tractable method for purification, a chemical trigger to reverse the stable imidazolinone ring in a traceless manner is needed. Thus, as described below, we herein report a method for the covalent, reversible, and efficient peptide capture and release^[22]^.

To understand substituent effects on the peptide capture we screened aromatic and heteroaromatic aldehydes possessing different rings, heteroatoms, and regiochemical placement of the aldehyde. In total 30 aldehydes were tested and ranked in order of the amount of N-terminally capped product, as an imidazolinone, formed on the model peptide Ser-Gly-Trp in a six-hour reaction at 37 °C in 50 mM sodium phosphate buffer pH 7.5 (Figure 1). Each reaction was analyzed by LC/MS, and aminal formation was confirmed by the presence of two distinct peaks in the 218 nm HPLC trace which had masses corresponding to the imidazolinone capped product. The two peaks arise from the two diastereomers formed during the ring closure.

**Figure 1.**
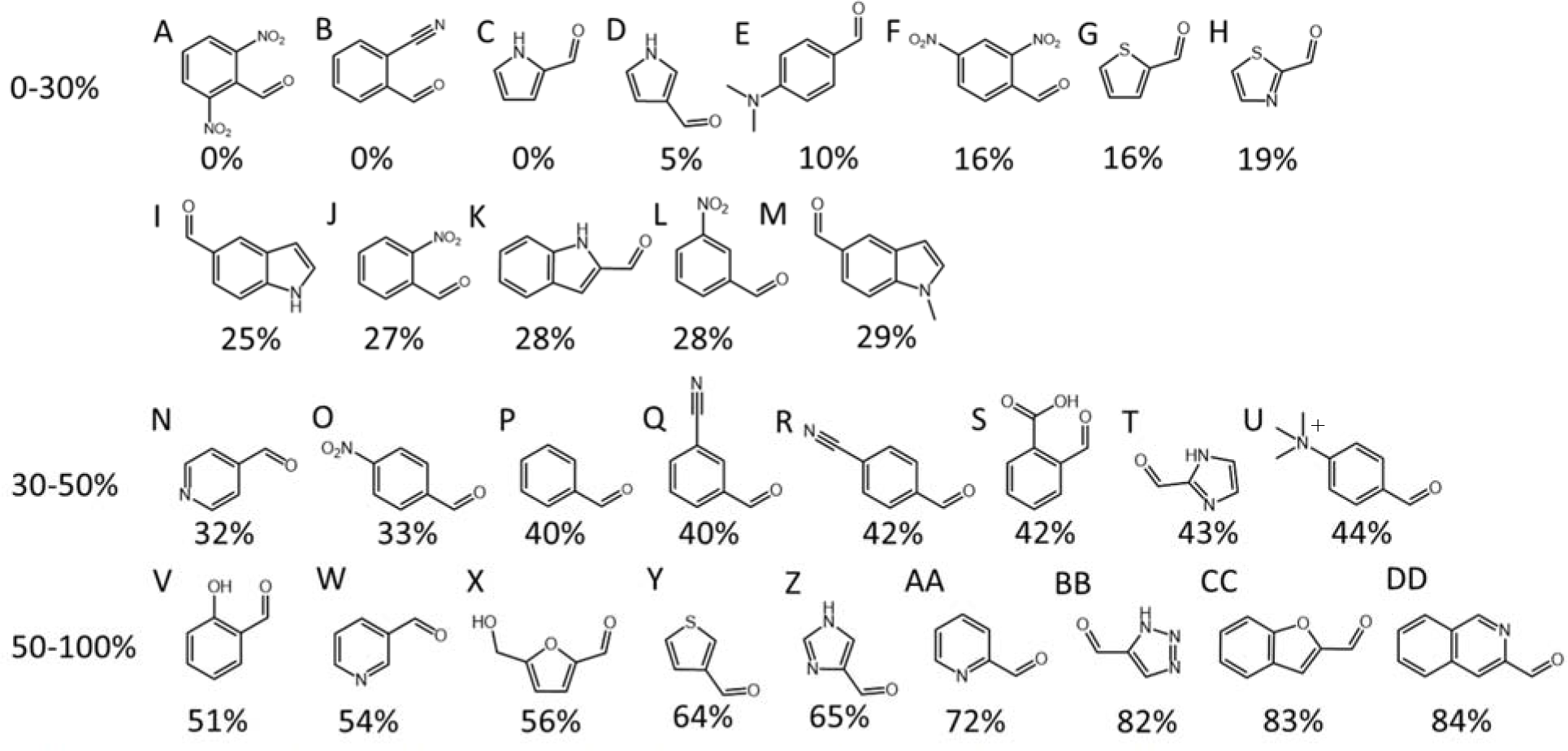
A-DD) are the aldehydes screened to find compounds that undergo aminal formation with the N-terminus of a peptide, without derivatization of lysine. Each shows the structure and the percent aminal formed based on the quantitation of the area under the curve from the HPLC of the reaction. Reaction conditions: 1 mM Ser-Gly-Trp peptide in 50 mM sodium phosphate buffer pH 7.5 is mixed with each aldehyde (4 mM final concentration) solubilized in DMF. These are shaken at 37 °C for 6 hours prior to LC-MS analysis (Buffer A: H_2_O + 0.1 % formic acid; Buffer B: methanol + 0.1 % formic acid). Each reaction was performed in triplicate.

We found that compounds that contain strongly electron-withdrawing groups (Fig 1. A and F) do not lead to imine-intermediates, the step required prior to ring closure. We postulate that this is due to the aldehyde being largely hydrated, which does not reverse to allow imine formation. However, less electron withdrawing character facilitates some product formation, but remains at unacceptable yields (J, L, N, O, Q, R, W). Imidazolinone formation is also disfavored when the aldehyde is on an electron rich aromatic ring, such as a thiazole/pyrrole (C, D, E, G, H, K), or has a substituent with a large negative Hammett sigma-value (M, albeit a Hammett plot was not generated). Aldehydes that are known to promote the formation of the imine complex through intramolecular hydrogen bonding or through a general-acid catalyzed mechanism^[23]^, albeit having a negative Hammett value, can promote product formation (V).

Clearly some electronic withdrawing character is necessary to facilitate nucleophilic attack of the N-terminal amine and ring closure with the adjacent amide, but not so much as to favor hydration. Thus, it appears that electron withdrawing heteroatoms adjacent to the aldehydes, exemplified as pyridines, triazoles, imidazoles, and furans (Z, AA, BB, CC, and DD) promote the imidazolinone formation.

The five most reactive aldehydes were selected to further test the ability for this reaction to discriminate the N-terminus of a peptide of the amine from an internal Lys residue. For this, we used a H_2_N-Ser-Gly-Lys-Trp-COOH peptide that was solubilized at 1 mM in 50 mM sodium phosphate buffer pH 7.5 and incubated with the aldehydes (4 mM final concentration) at 37 °C for six hours. The reactions were analyzed by LC/MS. The only detectable masses were two characteristic diasteromeric peaks that are present when the imidazolinone product is formed. Further, the five aldehydes showed similar imidazolinone formation as in the initial screen, and no product could be detected that corresponded to peptide with both an N-terminal imidazolinone and an imine on the lysine side chain (Figure 2). From the lack of product containing 2 aldehyde attached products, we confirm that these five aldehydes are N-terminal specific.

**Figure 2.**
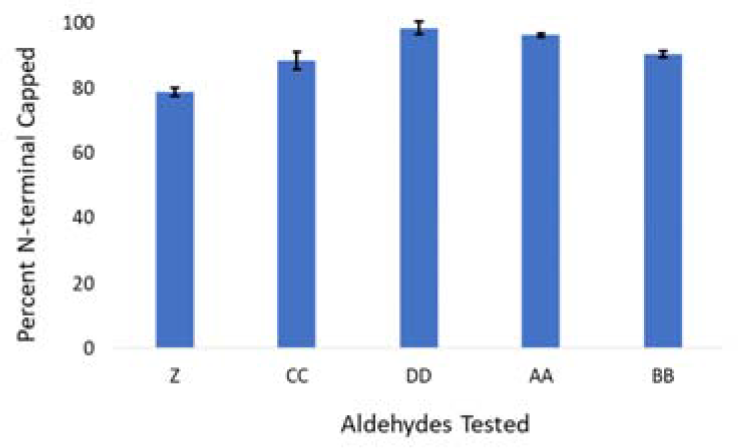
Percent of N-terminal capped product of SGKW peptide (4 eq) with aldehydes (1eq). Labels are taken from Figure 1. Z) 4-imidazolecarboxaldehyde AA) 2-pyridinylcarboxaldehyde BB) 1H-1,2,3-Triazole-5-carbaldehyde CC) benzofuran-2-carboxaldehyde DD) 3-formylisoquinoline.

Next, we moved on to use the N-terminal specific capture of peptides as a method for the rapid preparation of samples for traditional mass spectrometry-based proteomics. To do this, we used the water swellable PEG amine resin and performed amide bond formation to generate a resin that contained an acid cleavable Rink linker to which we added 6-formyl picolinic acid (FPCA) (Figure 3). A PCA derivative was used over one of the three derivatives found to perform more efficient capture as FPCA is available commercially and this will ensure this method can be used by the proteomics community more easily.

**Figure 3.**
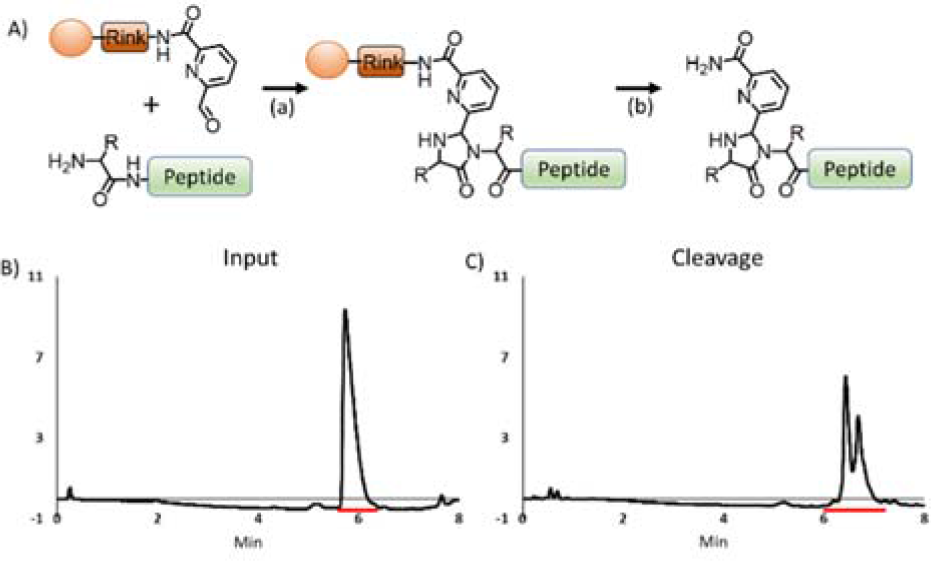
A) Schematic showing the PEG-Rink-FPCA resin synthesized and the steps for coupling and releasing peptides: (a) peptide in 50 mM sodium phosphate pH 7.5 is added to the resin and incubated at 37 °C for 16 hours, (b) the peptide is liberated from the resin using 95% trifluoroacetic acid, 2.5% H_2_O, and 2.5% triisopropyl silane for 2.5 hours B-C) HPLC of angiotensin I input (280 nm trace) (A) and TFA cleavage (B) after capture on PEG-Rink-FPCA resin. Red line indicates area under curve used to quantitate percent of peptide captured.

To evaluate the extent of capture and release by the aldehyde resin, we used angiotensin I as a test peptide. Capture of the peptide was determined by comparing the integrated peaks corresponding to the peptide during RP-HPLC analysis of the initial solution and the peptide resulting from the TFA cleavage. The captured peptide was released from the resin using a TFA cocktail (95 % TFA, 2.5 % H_2_O, 2.5% triisopropylsilane) to free the capped peptide. The resulting pellet was resuspended and analyzed with high-resolution mass spectrometry, and by comparing the input and cleavage sample we see an overall 70% yield of peptide. The capped peptide exists as diastereomers and the two peaks were found in the chromatogram (Figure 3 C).

With a highly efficient capture resin in hand, we then turned to using this as a method for the rapid purification and analysis of proteins from cell lysates for mass spectrometry-based proteomics. We lysed human embryonic kidney (HEK293T) cells by sonication. After isolating the soluble protein fraction, we denatured the proteins using trifluoroethanol and treated with iodoacetamide to alkylate the Cys residues. Next, dithiothreitol was added, and this mixture was incubated for 16 hours at 37 °C with PEGA-PCA resin and N-methylated trypsin in 50 mM phosphate buffer pH 7.5. This allowed for simultaneous proteolysis and capture of the resulting tryptic peptides. Afterwards the resin was washed extensively, and this step removed the small molecules (iodoacetamide and DTT), salts (phosphate buffer), and the trypsin. Finally, the peptides were released using the TFA cocktail, precipitated, and analyzed by high-resolution tandem mass spectrometry to identify the peptides and their parent protein.

Mass spectrometry analysis of the released peptides identified roughly 40,000 bead-bound peptides per replicate. More than half of the peptides observed did not possess a PCA adduct, attributed to non-specific adsorption to the resin as is commonly seen in resin-based purification from total cell lysates. Irrespective, from this data we were able to analyze for inherent biases in which peptides from the complex mixture bound specifically through their N-termini. We observed only a minor bias (~2.5x) for Ala at the N-terminal position and a minor bias (~3x) against Met (Figure 4B). Additionally, peptides with 2^nd^ position proline do not react with PCA and are selected against in this method.

**Figure 4.**
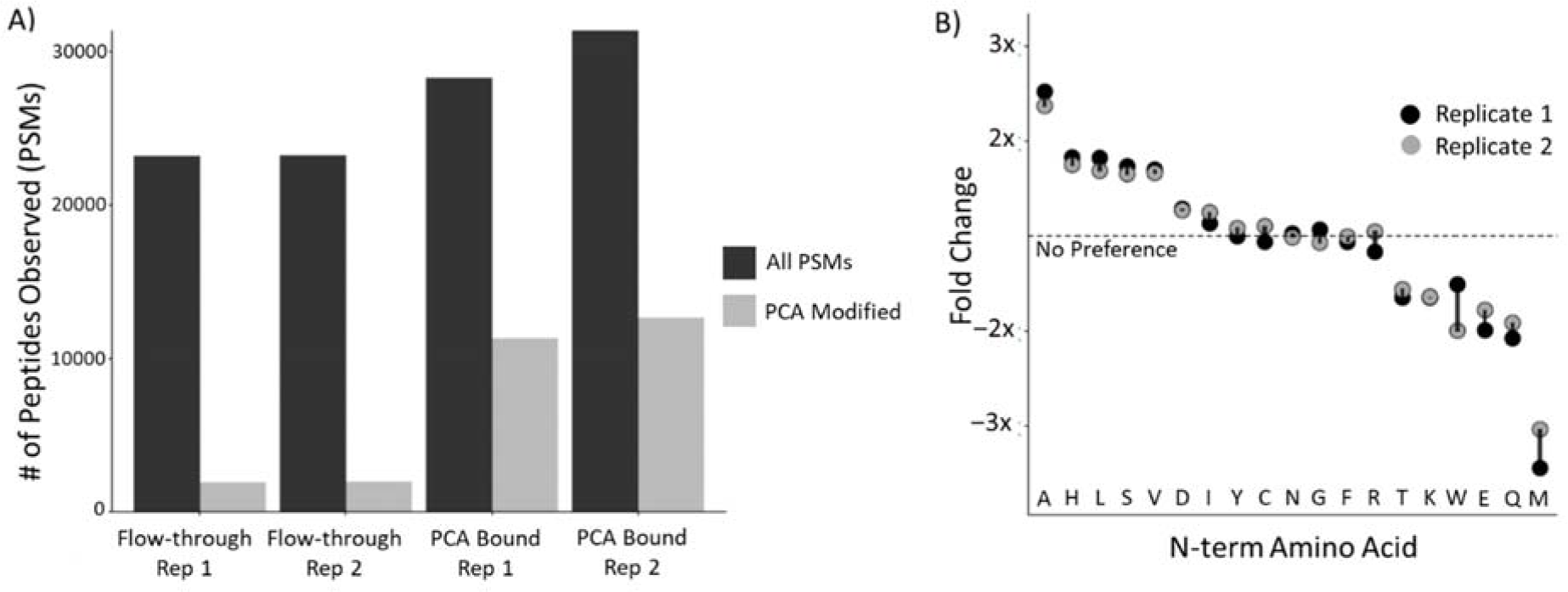
A) The number of unique proteins identified with high confidence (FDR < 1 %) in two replicate HEK293T cell lysate proteomics experiments. B) The normalized bias of the PCA capture reagent against the N-termini of the peptides bound to the resin. A total of 133,793 tandem mass spectra were collected and assigned, corresponding to 39,581 (from the bound sample) and 25,049 (from the flowthrough) in replicate 1, and 43,986 (from bound) and 25,177 (from flowthrough) in replicate 2.

The highly efficient peptide capture exhibited by the resin suggested it might prove useful for proteomics of low abundance samples, in particular by allowing for more effective fluorescent labeling of peptides in preparation for single molecule sequencing using a new proteomics technique known as fluorosequencing^[24]^. For this approach, peptides with fluorescently labeled amino acids are covalently tethered to a microscope cover slip and imaged at the single-molecule level using total internal reflection fluorescence microscopy (TIRF) following consecutive removals of N-terminal amino acids by Edman degradation. Edman cycles in which fluorescently labeled amino acids are removed reveal the amino acid sequence positions of the labeled amino acids, thus providing a peptide fingerprint on a molecule-by-molecule basis.

**Scheme 2.**
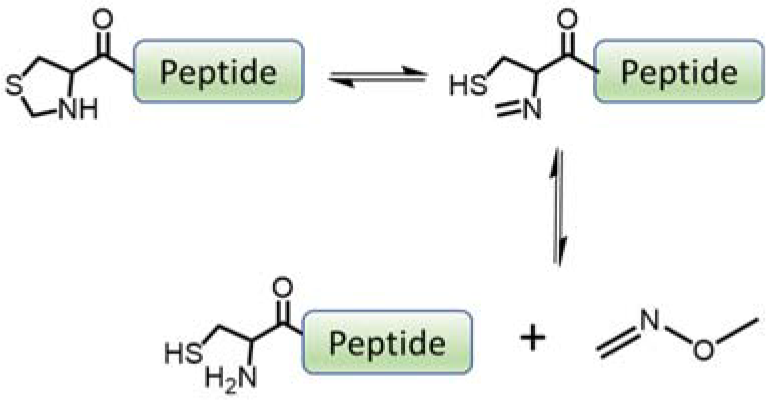
Deprotection of a thiazolidine peptide with methoxyamine.

In order for the FPCA resin to be useful for fluorosequencing, the N-terminus of the peptide must be protected during the initial labeling of amino acids to minimize side reactions when Lys is labeled. Another requirement is that the N-terminus must be free to allow for Edman degradation. Very recently, the Francis lab showed that when peptides attached to a resin bound PCA are treated with hydroxylamine the peptides slowly are released into solution over 6-10 days^[25]^. Concurrently, we had also optimized a similar reaction and found it was amenable to rapid analysis of peptides. To do this, we took inspiration from the field of Native Chemical Ligation^[26]^ and its use of thiazolidine, a protected variant of Cys, where a formaldehyde is condensed with the N-terminus and side chain to form a 5-membered ring (Scheme 2).

To reveal the Cys, it is common to utilize 300-400 mM methoxyamine at 37-42 °C, which quantitatively deprotects Cys in 3-4 hours^[27]^. With this as a starting point, we purified peptides that were N-terminally capped with either 4-nitrobenzaldehyde, PCA, or 3-formylisoquinoline. The status of the N-terminal cap was determined by ^1^H-NMR (Supplemental Figure 2-4) and no presence of the imine proton could be detected. Next, the peptides were treated with 300 mM methoxyamine at pH 3 and 60 °C for 24 hours (Figure 5 B). We observed a range of N-terminal deprotection levels, ranging from ~45% - 70% for the three different aldehydes tested. To optimize this reaction for speed and quantitative conversion, we utilized the more reactive dimethylaminoethyl hydrazine (DMAEH), which is extremely rapid in the formation of hydrazones^[28]^. After a 24 hr treatment of 300 mM DMAEH at pH 3 and 60 °C we observed nearly quantitative deprotection for all three aldehydes (Figure 5 C). This deprotection condition was used for future experiments.

**Figure 5.**
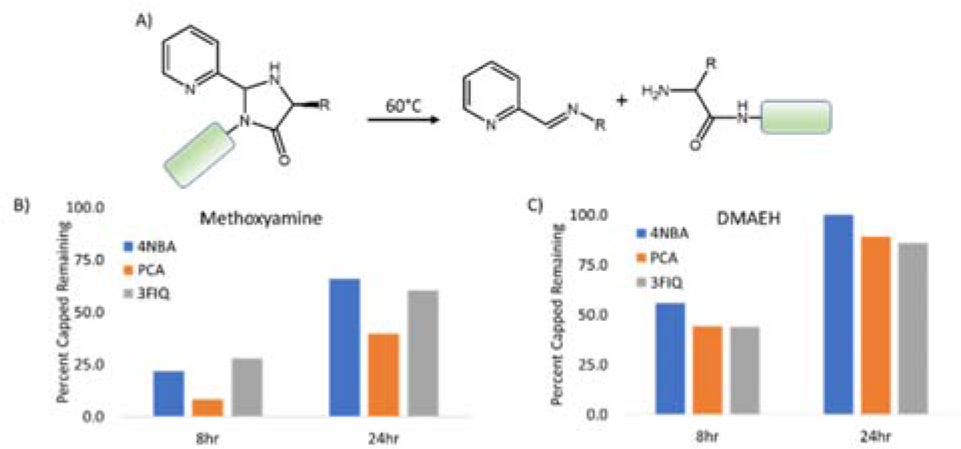
Reversal tests of N-terminally imidazolinone capped SGW peptide with 4-nitrobenzaldehyde (4NBA), 2PCA, or FIQ using either 0.3 M methoxyamine or 0.3 M dimethylaminoethyl hydrazine.

With a working reversal condition, we turned to the preparation of fluorescently labeled peptides. We considered a scenario in which peptides would be captured by their N-termini on resin, their C-termini modified with an alkynyl linker, one or more of their internal amino acids fluorescently labeled, and then peptides released. This scenario would test a number of important chemical steps and the ability to remove free fluorescent dyes to a level they cannot be detected by single molecule proteomics. To test if such a strategy was feasible with the PCA resin, we captured a short peptide (H_2_N-Ala-Lys-Ala-Gly-Ala-Gly-Arg-Tyr-Gly-COOH) onto the resin. Note that this peptide lacks acidic residues to avoid double labeling them with the C-terminus. Peptides from native sources would require C-terminal differentiation or be generated by a protease such as GluC whose action leaves all acidic residues positioned at peptide C-termini. We labeled the C-terminus with propargyl amine using HCTU/DIEA chemistry in DMF. Next, the Lys was labeled using Atto647N NHS esters in DMF. Finally, the peptide was cleaved from the resin using the TFA cocktail, precipitated, and the pellet was allowed to air dry.

Fresh azide-silane slides were prepared suitable for fluorosequencing, and the fluorescent peptide attached to their surface by Cu(I)-Click (Figure 6 A). The slide was washed extensively in ddH_2_O, and the PCA deprotected by submerging the slide in a bath of 500 mM DMAEH for 16 hours at 60 °C. Next, the slide was washed in water and assembled into the flow cell of the TIRF microscope and subjected to fluorosequencing as previously described^[24]^. We observed a strong loss of fluorescence after the second Edman cycle in 50% of the molecules, indicating that the Lys-Atto647N was removed at the appropriate Edman cycle as expected (Figure 6 B-D). The remaining peptide losses are accounted by errors produced during the physico-chemical processes such as photobleaching, Edman inefficiency etc, and are similar to the results obtained with peptides generated on the slides from Fmoc-N-terminal protection. These errors have been previously described and modeled^[24]^. Thus, this capture/label/release strategy offers a novel and elegant solution for proteomics experiments.

**Figure 6.**
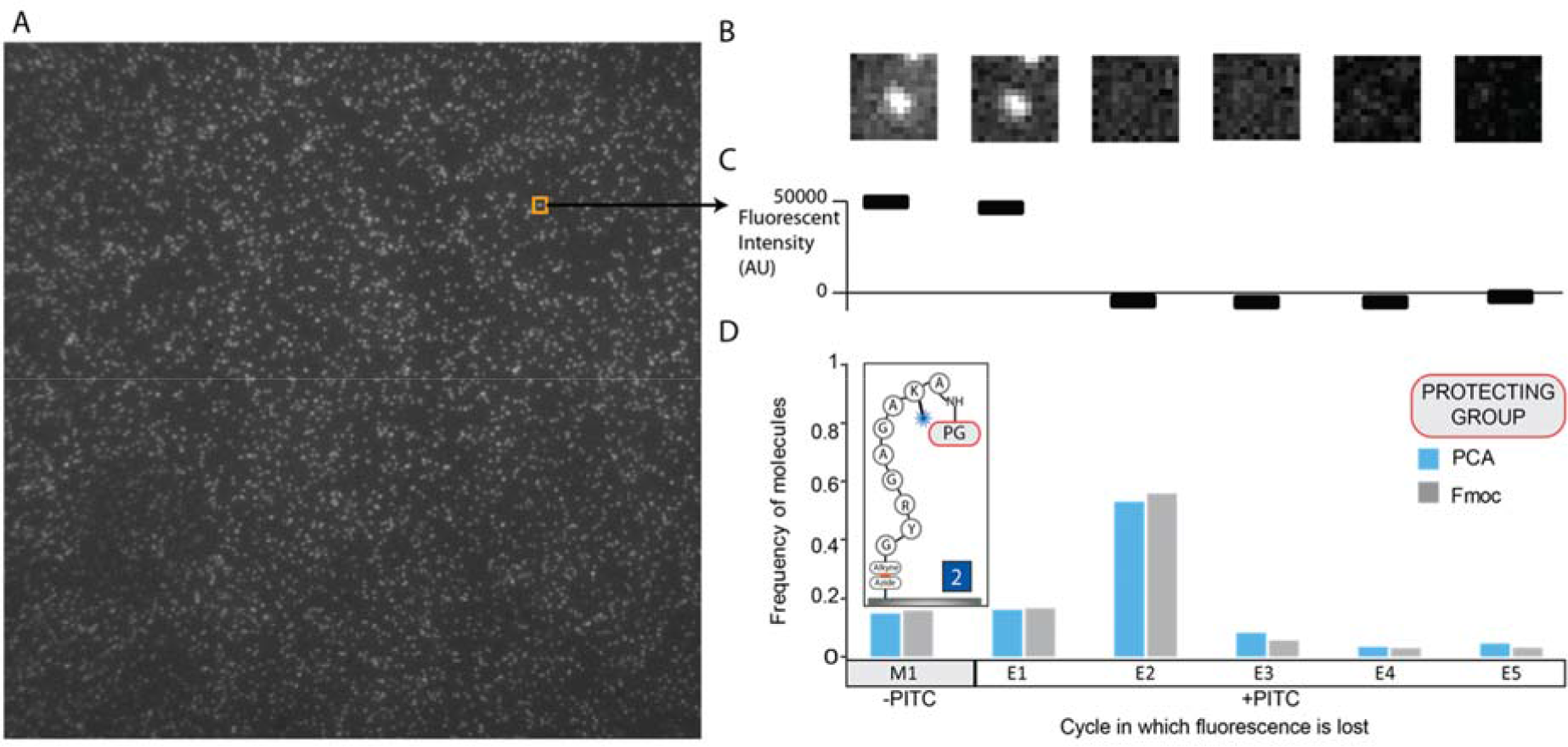
A) Representative field of view from a fluorosequencing experiment. B) Tracking an individual peptide across the Edman cycles with its subsequent loss of fluorescence after cycle 2. C) Fluorescent intensity of the peptide measured across Edman cycles. D) Frequency of molecules lost after each step for both a PCA or Fmoc protected peptide. M1 is a "mock" cycle where the slide is washed with all reagents used in fluorosequencing without the phenyl isothiocyantate (PITC) being present to remove non-specifically bound peptides or free fluorophores.

## Conclusion

Here, we have shown that the ability to covalently bind peptides from a complex mixture on a solid phase substrate, anchoring them by their N-termini and allowing for their efficient purification for multiple types of proteomics experiments. Further, we show this method allows for the removal of salts, small molecules and even the proteases used to prepare samples. This methodology will also allow removal of any other non-peptide contaminant in the sample including detergents.

Importantly, binding the peptides to the resin allows for the use of a number of organic solvents, as the PEG resin is compatible with many solvents that peptides generally are not fully soluble in. We utilized this feature to chemically modify the C-termini of peptides bound to the resin using standard peptide synthesis techniques and fluorescently label Lys residues of bound peptides using an NHS ester in DMF and organic bases.

For experimental techniques that require multiple chemical reactions such as fluorosequencing or nanopore sequencing^[29]^, we expect the resin will prove valuable for the analysis of proteomic samples with small sample sizes. Working with smaller protein concentrations is an important step for these single-molecule techniques, and advances proteomic techniques to be able to analyze clinically relevant samples, such as tumor biopsies.

## Experimental Section

### Materials

Chemicals were purchased and used as received without any purification. 6-formyl picolinic acid was purchased from Enamine. Atto647N-NHS was purchased from Attotek. Azide functionalized slides were purchased from PolyAn. All other chemicals were purchased from Sigma Aldrich.

### Peptide Synthesis

All peptides were synthesized using automated microwave-assisted solid-phase peptide synthesis (Liberty Blue Microwave Synthesizer, CEM Corporation). Synthesis was performed using standard Fmoc chemistry using DIC/Oxyma coupling strategies (1:1:1 ratio with amino acids). Coupling steps were performed at 90 °C for 120 seconds, and deprotection was performed using 20 % piperidine in DMF at 90 °C for 60 seconds. All peptides were cleaved from resin using trifluoroacetic acid (TFA), triisopropylsilane (TIS), and H_2_O (95:2.5:2.5) for 2.5 hours prior to the cleavage solution being concentrated under nitrogen stream. The peptide is precipitated with ice cold diethyl ether and collected by centrifugation at 12,000 *g* for 10 minutes. Peptides were purified using a Grace-Vydac C18 column (Buffer A: H_2_O + 0.1 % formic acid; Buffer B: methanol + 0. 1% formic acid) over a 10-60 % gradient.

### In-solution N-terminal Peptide Capture

Peptides were mixed with four molar equivalents of aldehyde in 50 mM phosphate buffer pH 7.5, then incubated for 8-16 hours at 37 °C prior to purification or analysis. All aldehydes used were solubilized in DMF at 100 mM then diluted to final concentration. Samples were analysed by LC/MS. Aminal formation was determined by quantitation of remaining unreacted peptide using HPLC.

### Reversal of Aminal Cap

Peptides were first allowed to react with 4-nitrobenzaldehyde, 2-pyridinylcarboxaldehyde, or 3-formylisoquinoline following standard in-solution reaction procedures (4 mM aldehyde and 1 mM peptide). Peptides were then purified using a Grace-Vydac C18 RP-HPLC column, analysed by LC/MS and lyophilized to dryness. Complete imidazolinone conversion was checked by ^1^H-NMR (MR400 MHz, Agilent) For the reversal tests, capped peptides were resuspended in either 0.3 M dimethylaminoethyl hydrazine or 0.3 M methoxyamine. Samples were incubated at 60 °C and then analysed by HPLC and mass spectrometry at each time point. Percentage released is determined by comparing the integration of the HPLC peak of the capped peptide over time.

### Aldehyde Capture Resin Preparation

Amino PEGA resin (Novabiochem) was used and was functionalized with Fmoc-Peg_2_-OH, Rink linker and 6-formylpyridine-2-carboxylic acid using HCTU/DIEA (1:1:1.1 ratio) chemistry coupling for 45 minutes. Deprotection was done using 20 % piperidine in DMF two times for five minutes each. Resin was stored in DMF at 4 °C prior to use.

### Resin-based Peptide Capture

Capture resin was washed in DMF, water, and 50 mM phosphate buffer pH 7.5. Each wash included a 5 minute incubation in the solvent. Peptide was then added to the resin in 50 mM phosphate buffer pH 7.5 and incubated at 37 °C for 16-24 hours. Next the resin was washed extensively in incubation buffer, water, and finally DMF. After derivatization, the resin was washed extensively in water, DMF, and finally DCM. The peptide was cleaved from resin in 95 % TFA, 2.5 % TIS, and 2.5 % H_2_O. The TFA was concentrated under N_2_ stream and ether precipitated prior to mass spectrometry analysis.

### Cell Growth Conditions

HEK-293T cells were grown in Dulbecco’s Modified Eagle Medium with 10 % Fetal Bovine Serum at 37 °C and 5 % CO_2_. Cells were passaged when between 70-80 % confluence.

### HEK Lysate Digestion and Capture

Cells were grown to 80 % confluence and harvested in PBS and pelleted at 500 *g* for 3 minutes. Cells then suspended in hypotonic 50 mM Tris-HCl buffer pH 8 and placed on ice. Protease inhibitor (Mini cOmplete, EDTA Free protease inhibitor cocktail, Roche) was added to 1x concentration. Cells were sonicated (Branson 2510) for 1 minute at 42 kHz and placed on ice for an additional minute. This was repeated 3 times. The solution was then centrifuged at 17,000 *g* for 10 minutes at 4 °C and the supernatant was collected. Protein content was then measured using a Bradford Assay. 250 µg of protein was denatured in 2,2,2-trifluoroethanol (TFE) and 5 mM tris(2-carboxyethyl)phosphine (TCEP) at 45 °C for 45 minutes. Proteins were alkylated in the dark with 5.5 mM iodoacetamide, then the remaining iodoacetamide quenched in 100 mM dithiothreitol. MS-grade reductively methylated trypsin (Pierce) was then added to the solution in a ratio of 1:25. This mixture was added to PEGA-FPCA beads and diluted with 100 mM phosphate buffer pH 7.5. The peptide mixture was incubated at 37 °C for 18 hours. Next, the resin was washed extensively with H_2_O, DMF and DCM prior to cleaving the resin with a TFA cocktail (95 % trifluoroacetic acid, 2.5 % triisopropylsilane, and 2.5 % H_2_O) for 2.5 hours. The peptides were precipitated with ice cold ether and allowed to air dry.

### Mass Spectrometry

Peptides were separated on a 75 µM x 25 cm Acclaim PepMap100 C-18 column (Thermo Scientific) using a 3-45 % acetonitrile + 0.1 % formic acid gradient over 120 min and analysed online by nanoelectrospray-ionization tandem mass spectrometry on an Orbitrap Fusion (Thermo Scientific). Data-dependent acquisition was activated, with parent ion (MS1) scans collected at high resolution (120,000). Ions with charge 1 were selected for collision-induced dissociation fragmentation spectrum acquisition (MS2) in the ion trap, using a Top Speed acquisition time of 3-s. Dynamic exclusion was activated, with a 60-s exclusion time for ions selected more than once. MS proteomics data were acquired in the UT Austin Proteomics Facility and have been deposited to the ProteomeXchange Consortium *via* the PRIDE partner repository with dataset identifier PXD016291 and 10.6019/PXD016291.

### Protein Identification

Proteins were identified with Proteome Discoverer 2.3 (Thermo Scientific) against the Uniprot human reference proteome. To distinguish PCA protected peptides, searches incorporated an optional peptide N-terminal dynamic modification (132.032 Da) corresponding to the PCA modified peptide, thresholding identifications with a false discovery rate of 1 %.

### On Bead Labeling of Peptides

Peptides were captured to PCA resin as described. After rinsing the C-terminus was first coupled with 100 mM propargylamine, 100 mM HCTU, and 100 mM DIEA in DMF for 2 hours at room temperature. The resin was extensively washed with DMF and the Lys residues were labelled with 0.5 mM Atto647N-NHS (Attotec) and 1 mM DIEA overnight at room temperature. The resin was washed extensively in DMF and DCM and all of the peptides cleaved from the resin with a TFA cocktail (95 % TFA, 2.5 % H_2_O, and 2.5 % TIS) for 2.5 hours. The supernatant was collected and concentrated with a N_2_ stream. Ice cold diethyl ether is added (10 vol) and the peptides collected by centrifugation for 10 minutes at 17,000 *g*. The peptide was analysed by high-resolution mass spectrometry to confirm the double labelling.

### Single Molecule Peptide

Sequencing: Approximately 200 pM of peptides are immobilized on an azide slide (custom slides from PolyAn, Germany) using standard Cu(I)-Click chemistry. Briefly a 2 mL solution comprising peptide (200 pM), CuSO_4_/tris-hydroxypropyltriazolylmethylamine (THPTA) mix (1 mM/0.5 mM) and freshly prepared sodium L-ascorbate (5 mM) was incubated on the azide slide at room temperature for 2 hours. Following the incubation, the slides were rinsed with water and fluorosequencing performed as previously described with minor modifications^[24]^. To deprotect the N-terminal PCA cap, the slides were bathed in 0.5 M DMAEH at 60 °C for 16 hours. Deprotection of the Fmoc group was performed by incubating slides with 20% Piperidine solution in DMF for 1 hour. The images were processed using custom developed script (available at https://github.com/marcottelab/FluorosequencingImageAnalysis/).

## Acknowledgements

This work was supported by grants from DARPA (N66001-14-2-4051 to EVA and EMM), NIH (R35 GM122480, R01 HD085901, R01 DK110520 to EMM), Erisyon, Inc. (to EMM and EVA), and the Welch foundation (F-1515 to EMM and F-0046 to EVA). JS, EMM, AB and EVA are co-founders and shareholders of Erisyon, Inc. A patent application based on aspects of this work is pending.

