## Supplemental Data for "Solid-Phase Peptide Capture and Release for Bulk and Single-Molecule Proteomics"

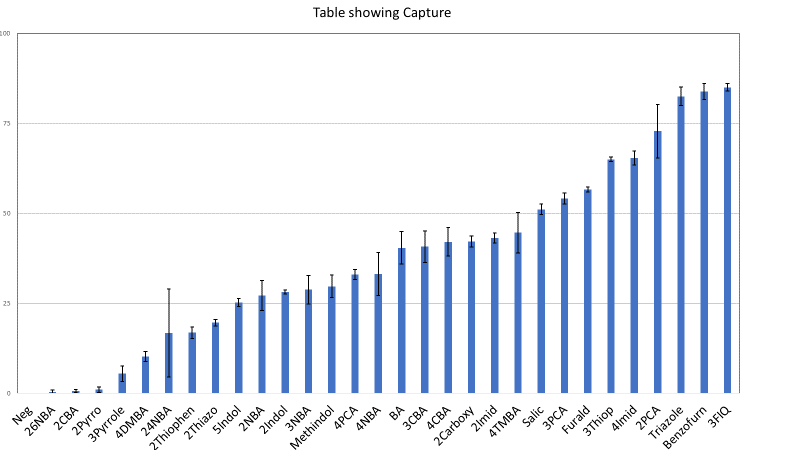

**Figure 1:** **Graphic representation of the aldehydes screened**

Aldehydes tested in Main Figure 1. Bars indicate percent peptide capped by PCA. Error bars are the standard deviation based off of three replicates: A) 2,6-dinitrobenzaldehyde B) 2-cyanobenzaldehyde C) Pyrrole-2-carboxaldehyde D) Pyrrole-3-carboxaldehyde E) 4-(dimethylamino)benzaldehyde F) 2,6-dinitrobenzaldehyde G) Thiophene-2-carboxaldehyde H) 2-Thiazolecarboxaldehyde I) Indole-5-carboxaldehyde J) 2-nitrobenzaldehyde K) Indole-2-carboxaldehyde L) 3-nitrobenzaldehyde M) 1-methyl-1H-indole-5-carboxaldehyde N) 4-pyridinylcarboxaldehyde O) 4-nitrobenzaldehyde P) benzaldehyde Q) 3-cyanobenzaldehyde R) 4-cyanobenzaldehyde S) 2-carboxybenzaldehyde T) 2-imidazolecarboxaldehyde U) 4-(trimethylamine)benzaldehyde V) salicylaldehyde W) 3-pyridinecarboxaldehyde X) 5-hydroxymethyl-2-furaldehyde Y) thiophene-3-carboxaldehyde Z) 4-imidazolecarboxaldehyde AA) 2-pyridinylcarboxaldehyde BB) 1H-1,2,3-Triazole-5-carbaldehyde CC) benzofuran-2-carboxaldehyde DD) 3-formylisoquinoline.

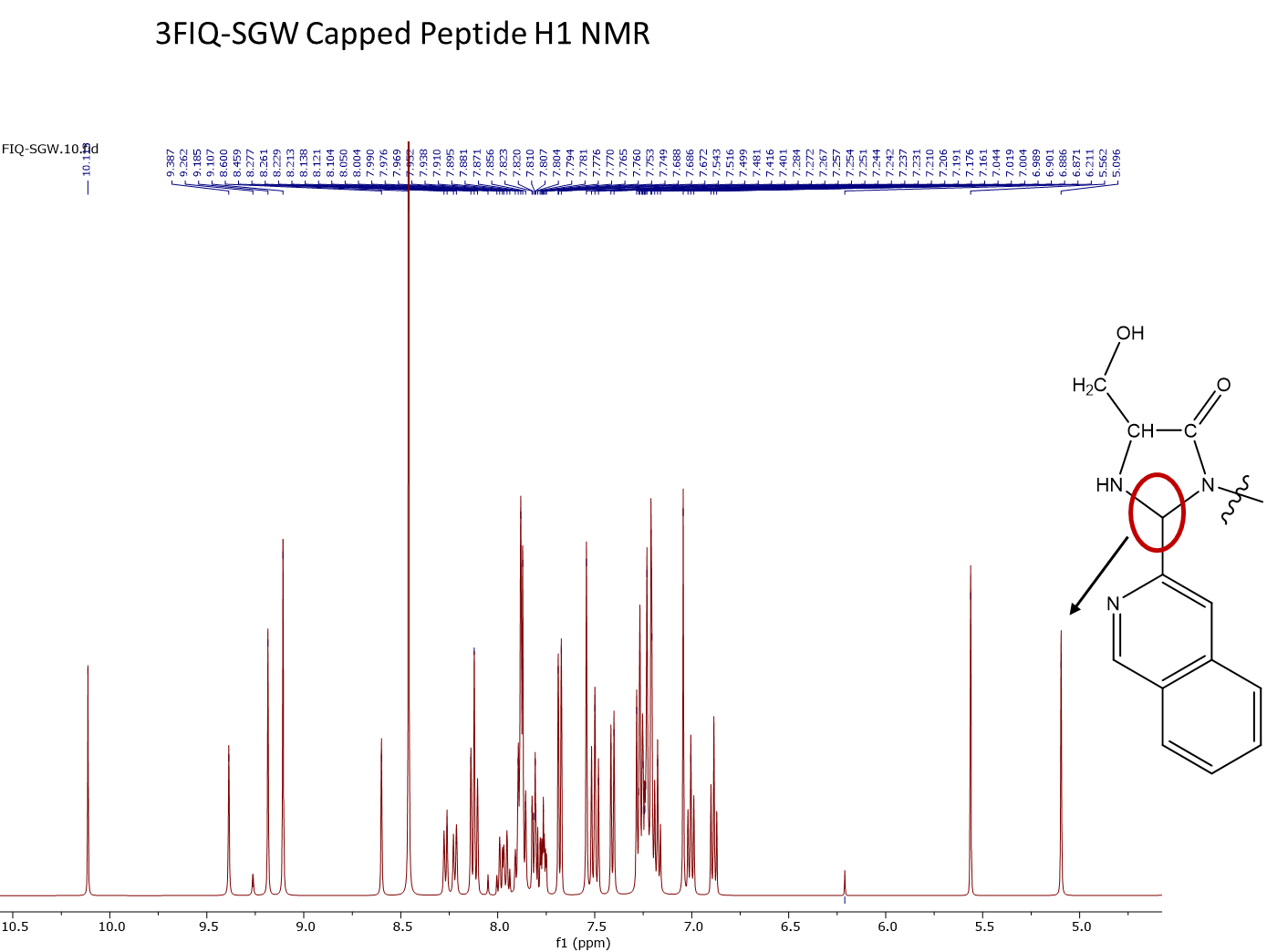

**Figure 2: ^1^H-NMR of 3-formylisoquinoline-SGW peptide.**

^1^H-NMR was taken in D_2_O of purified aldehyde labeled Ser-Gly-Trp peptide with the imidazolinone proton labeled. No proton that corresponded to the aldehyde or imine was observed

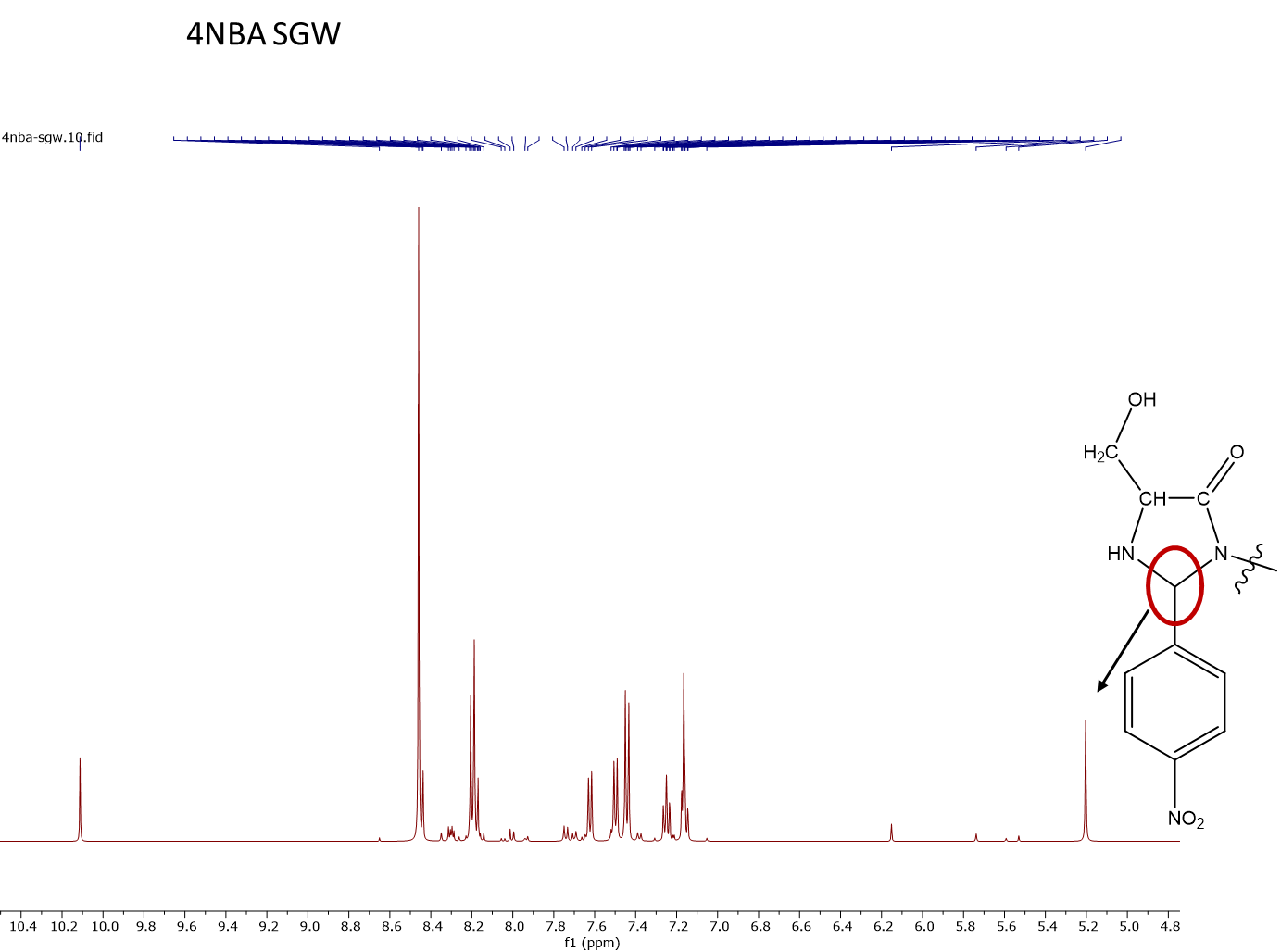

**Figure 3: ^1^H-NMR of 4-nitrobenzaldehyde-SGW peptides.**

^1^H-NMR was taken in D_2_O of purified aldehyde labeled Ser-Gly-Trp peptide with the imidazolinone proton labeled. No proton that corresponded to the aldehyde or imine was observed

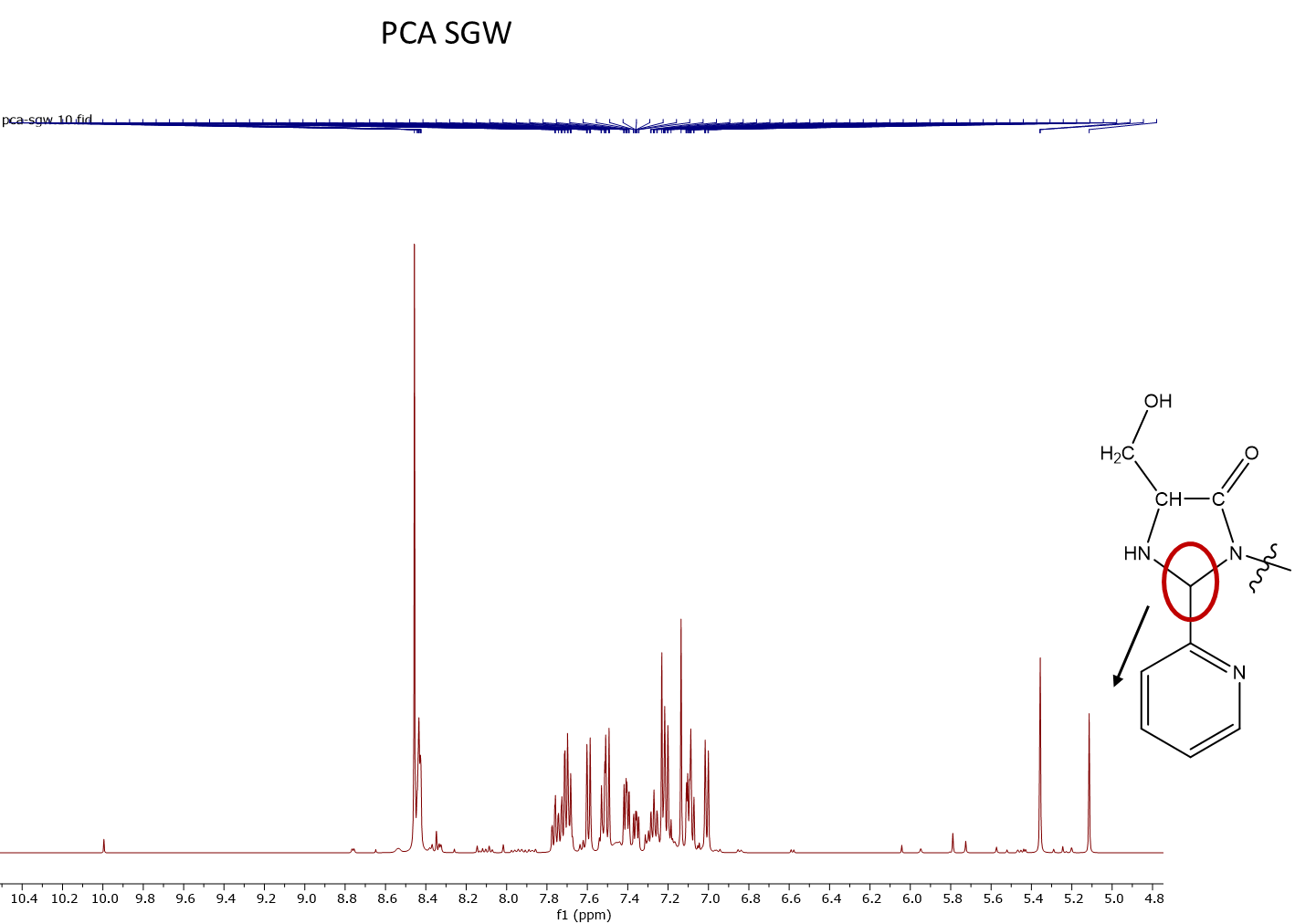

**Figure 4: ^1^H-NMR of pyridinylcarboxaldehyde-SGW peptide.**

^1^H-NMR was taken in D_2_O of purified aldehyde labeled Ser-Gly-Trp peptide with the imidazolinone proton labeled. No proton that corresponded to the aldehyde or imine was observed

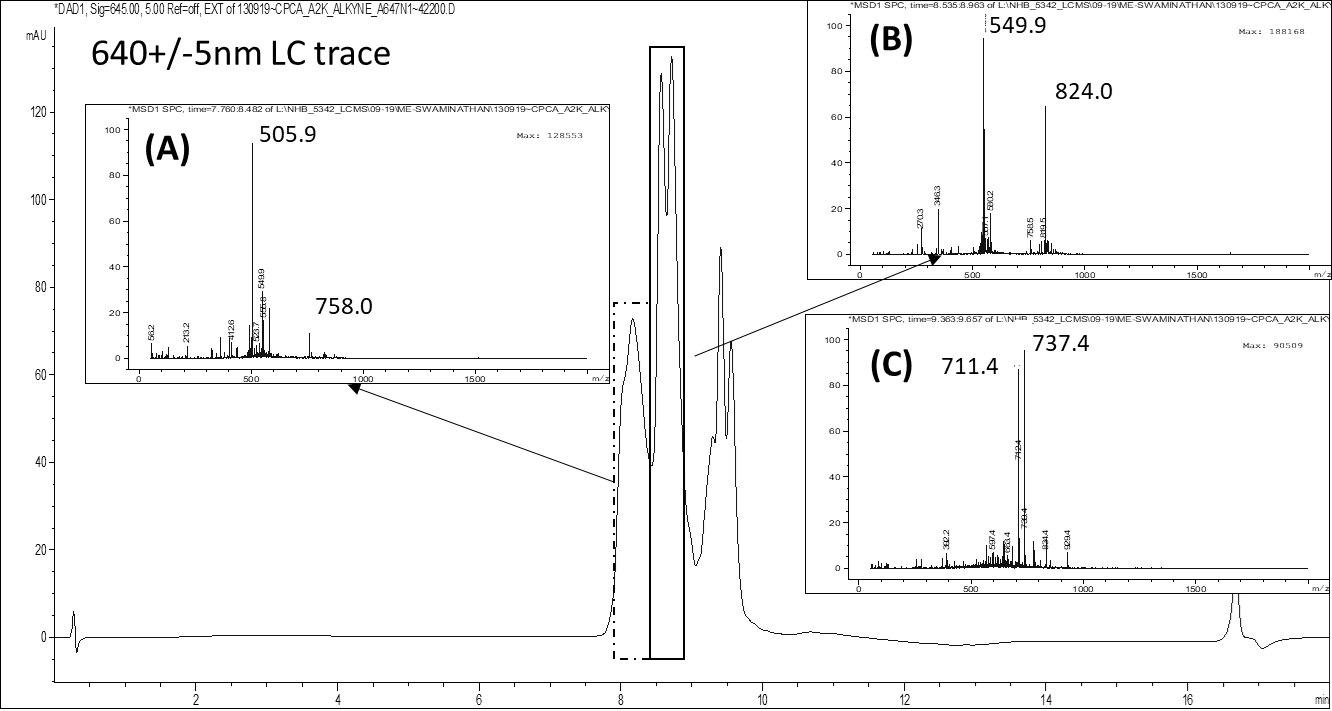

**Figure 5: Multiple derivatizations on resin-captured peptides was performed and verified.**

The resin immobilized peptide’s (sequence H_2_N-AKAGAGRYG-OH) (1) C-terminal carboxylate was labeled with propargylamine and (2) amine side chain of lysine was labeled with Atto647N fluorophore. The 16 min gradient LC-MS analysis (see methods) indicated that >70% of the products observed with 640 nm LC trace (shown above) corresponds to the multiply labeled peptide. Insert A and B corresponds to the peptide with Atto647N dye and the alkyne label. While inset A corresponds to peptide without the N-terminal PCA adduct (Exp. Mass: [M+H]^+^ 1514.8 m/z, [M+2H]^+2^ 757.9 m/z), inset B is the PCA capped peptide (Exp. Mass: [M+H]^+^ 1646.9 m/z, [M+2H]^+2^ 823.9 m/z). Inset C indicates side products observed in the reaction

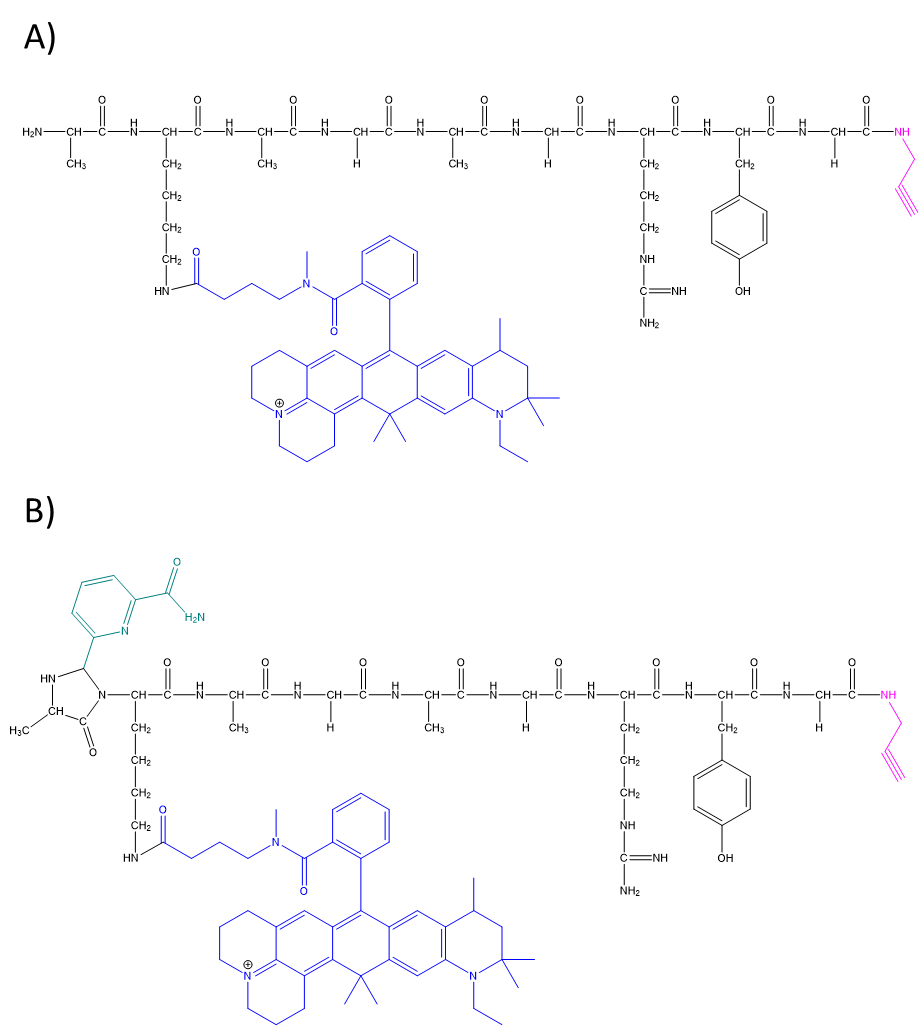

**Figure 6: Chemical structures** **of products from captured peptide labeling.**

Schematics of the peptides found in the LC/MS analysis that correspond to the molecular weights seen in Figure 5 A) The product in Figure 5 that is derivatized on the C-terminus and Lys but does not have an N-terminal PCA adduct. Exp. Mass: 1514.8 m/z B) The product in Figure 5 that has both modifications as well as an N-terminal adduct. Exp. Mass: 1646.9 m/z
